# Genome-wide association study of peripheral blood DNA methylation and conventional mammographic density measures

**DOI:** 10.1101/458299

**Authors:** Shuai Li, Pierre-Antoine Dugué, Laura Baglietto, Gianluca Severi, Ee Ming Wong, Tuong L. Nguyen, Jennifer Stone, Dallas R. English, Melissa C. Southey, Graham G. Giles, John L. Hopper, Roger L. Milne

## Abstract

Age- and body mass index (BMI)-adjusted mammographie density is one the strongest breast cancer risk factors. DNA methylation is a molecular mechanism that could underlie interindividual variation in mammographic density. We aimed to investigate the association between breast cancer risk-predicting mammographic density measures and blood DNA methylation. For 436 women from the Australian Mammographic Density Twins and Sisters Study and 591 women from the Melbourne Collaborative Cohort Study, mammographic density (dense area, non-dense area and percentage dense area) defined by the conventional brightness threshold was measured using the CUMULUS software, and peripheral blood DNA methylation was measured using the HumanMethylation450 (HM450) BeadChip assay. Associations between DNA methylation at >400,000 sites and mammographic density measures adjusted for age and BMI were assessed within each cohort and pooled using fixed-effect meta-analysis. Associations with methylation at genetic loci known to be associated with mammographic density were also examined. We found no genome-wide significant(*P*<10^−7^)association for any mammographic density measure from the meta-analysis, or from the cohort-specific analyses. None of the 299 methylation sites located at genetic loci associated with mammographic density was associated with any mammographic density measure after adjusting for multiple testing (all *P*>0.05/299 = 1.7 × 10^−4^). In summary, our study did not detect associations between blood DNA methylation, as measured by the HM450 assay, and conventional mammographic density measures that predict breast cancer risk.

## Introduction

Mammographie density has conventionally been defined as the white or bright region on a mammographic image of the breast. Mammographic density declines with increasing age and body mass index (BMI). After adjustment for age and BMI, mammographic density has been found to be one of the strongest risk factors for breast cancer in terms of differentiating cases from controls: age- and BMI-adjusted mammographic density has an odds per adjusted standard deviation (OPERA) of approximately 1.4, which is comparable to other strong breast cancer risk factors such as polygenic risk scores^1^. Women with more than 75% dense breast tissue have a four- to five-fold increased breast cancer risk compared with women of the same age and BMI with less than 10% dense breast tissue^2^.

Variation across the population in mammographic density has both environmental and genetic determinants. Age, BMI, and other breast cancer risk factors explain about 30% of the variation in mammographic density, and over 60% of the residual variation appears to be accounted for by additive genetic factors^3^. Genome-wide association studies (GWAS) have identified variants at specific genetic loci associated with mammographic density^4, 5^, and about 15% of the genetic variants known to be associated with breast cancer risk are also associated with mammographic density measures^6^, although they explain only a small fraction of the latter. Despite these known determinants, the molecular mechanisms underlying variation in mammographic density are not well understood, nor is how mammographic density translates into breast cancer risk at the biological level^7^.

DNA methylation, a process whereby typically a methyl group is added to a cytosine-guanine dinucleotide, plays an important role in modulating gene expression without changing the underlying DNA sequence. There is increasing evidence that blood DNA methylation is associated with variation in traits or disease risks, and also considerable interest in determining if there is any relationship between blood DNA methylation and breast cancer risk^8^. We tested the hypothesis that blood DNA methylation underlies inter-individual variation in breast cancer risk-predicting mammographic density measures using data for middle-aged women from two Australian cohorts.

## Material and methods

### Sample

The study sample included 436 women from the Australian Mammographic Density Twins and Sisters Study (AMDTSS) and 591 women from the Melbourne Collaborative Cohort Study (MCCS). Included participants from the two cohorts had, on average, similar characteristics (Table 1).

**Table 1.** Characteristics of samples from the two cohorts

| Sample characteristics | AMDTSS (N=436) | MCCS (N=591) |
| --- | --- | --- |
| Age at mammogram (years), mean ± SD | 55.6 ± 8.1 | 59.5 ± 7.3 |
| Age at blood draw (years), mean ± SD | 56.7 ± 7.9 | 57.2 ± 7.9 |
| Body mass index (kg/m<sup>2</sup>), mean ± SD | 26.8 ± 5.8 | 26.6 ± 4.5 |
| Post-menopausal, N (%) | 305 (70) | 435 (74) |
| Hormone replacement therapy use, N (%) | 53 (12) | 119 (20) |
| Number of live births, mean ± SD | 2.6 ± 1.5 | 2.4 ± 1.5 |
| Ever smoker, N (%) | 166 (38) | 198 (34) |
| Mammographic density measures |  |  |
| Dense area (cm<sup>2</sup>), median (IQR) | 25.6 (10.1-45.7) | 11.3 (4.2-20.7) |
| Non-dense area (cm<sup>2</sup>), median (IQR) | 99.1 (65.8-143.3) | 116.4 (82.7-159.2) |
| Percentage dense area (%), median (IQR) | 21.5 (7.0-39.6) | 9.5 (3.1-19.2) |

N, sample size; SD: standard deviation; IQR: inter-quartile range

The AMDTSS is a twin family cohort study of mammographic density^9^. Between 2004 and 2009, twins aged 40-70 years who participated in the Australian Twin Study of Mammographic Density between 1995 and 1999 were asked to participate further, and their non-twin sisters were also invited to participate. Participants completed questionnaires through telephone-administered interviews, and gave blood samples and permission to access their mammograms. A total of 479 women from 130 families oversampled for having one or more participating sisters were selected for DNA methylation research^10^. Among them, 436 women also had mammographic density data available and were included in this analysis. All women were of European descent.

The MCCS is a prospective cohort of 41,513 adults (24,469 women) aged between 27 and 76 years (99.3% aged 40-69) when recruited between 1990 and 1994^11^. Participants completed questionnaires in face-to-face interviews, had clinical measures taken by trained staff and gave a blood sample, the latter collected as dried blood spots on Guthrie cards, as peripheral blood mononuclear cells or as buffy coats. All MCCS participants were of European descent, including 25% born in Italy or Greece. Blood DNA methylation was measured for 6,404 participants in one of six nested-case control studies for breast cancer, colorectal cancer, mature B cell neoplasms, urothelial cell carcinoma, kidney cancer, or lung cancer^12, 13^. In the breast cancer study, only women who included as controls were selected, to avoid the possibility of collider bias^14^. A total of 591 women who had mammographic density and DNA methylation data available were included in this analysis.

### DNA methylation

DNA was extracted from peripheral blood samples and methylation was measured using the Illumina Infinium HumanMethylation450 (HM450) BeadChip and following the same protocol for both cohorts ^10, 13^. DNA was extracted from dried blood spots stored on Guthrie cards using a previously reported method^15^, and from peripheral blood mononuclear cells and buffy coat specimens using QIAamp mini spin columns (Qiagen, Hilden, Germany). Bisulfite conversion was performed using EZ DNA Methylation-Gold single-tube kit (Zymo Research, Irvine, CA) according to the manufacturer’s instructions. A total of 200ng of bisulfite-converted DNA was whole-genome amplified and hybridised onto HM450 BeadChips. DNA methylation was assessed according to manufacturer’s instructions. Raw intensity data was processed using the Bioconductor *minfi* package. Illumina’s reference factor-based normalization methods *(preprocessIllumina*) and subset-quantile within array normalization for type I and II probe bias correction^16^ were used for data normalisation. In the AMDTSS, the empirical Bayes batch-effects removal method *ComBat*^17^ was applied. Probes with missing value (detection *P*-value>0.01) in one or more samples were excluded. In the MCCS, samples were excluded if >5% probes had a detection P-value higher than 0.01, which were regarded as missing values, while probes were excluded from further analysis if more than 20% of samples had missing values. After quality control, 439,085 autosomal methylation sites were common to both datasets.

### Mammographic density measures

Both cohorts measured mammographic density using the same protocol^9, 18^. In the AMDTSS, mammograms were retrieved from BreastScreen Services, from private clinics, and from women who kept their films at home. In the MCCS, mammograms were retrieved from BreastScreen Victoria via record linkage. For a woman with several mammograms available,the mammogram taken closest to the date at blood draw was selected. Mammographic films were digitized by the Australian Mammographic Density Research Facility at the University of Melbourne. The digitized images were then masked so that extraneous features on the mammogram other than the breast were excluded, and randomized into reading sets of approximately 100 images. The total breast area and the dense area defined as the amount of the white or bright regions were measured by independent readers using a computer-assisted thresholding technique CUMULUS 3.0 (Imaging Research Program, Sunnybrook Health Sciences Centre, University of Toronto, Toronto, Canada). The non-dense area was calculated as the total breast area minus the dense area, and the percentage dense area was calculated as the dense area divided by the total breast area.

### Genome-wide association study

We used linear mixed-effects models to assess the association between the M-value (logit transformation of Beta-value, which quantifies the percentage of methylated cytosines) of each methylation site as the dependent variable, and each mammographic density risk measure as an independent variable. Because mammographic density measures were right-skewed, they were power-transformed to obtain approximately Gaussian distributions (dense area, power 0.25; non-dense area, power 0.4; percentage dense area, power 0.3). Models were adjusted by fitting fixed effects for age at blood draw (continuous), the difference between age at mammogram and age at blood draw (continuous), BMI (continuous), smoking status (never/ever in the AMDTSS and never/former/current in the MCCS), menopausal status (pre/post-menopausal), use of hormone replacement treatment (never/ever), number of live births (continuous), all collected at blood draw and cell type proportions (CD4T cells, CD8T cells, natural killer cells, B cells, monocytes, granulocytes) estimated from DNA methylation data using the Houseman algorithm^19^. For the AMDTSS, the model was additionally adjusted for family and zygosity as random effects. For the MCCS, the model was additionally adjusted for sample type (dried blood spots/mononuclear cells/buffy coats) and country of birth (Australia/UK/Italy/Greece) as fixed effects, and for study, plate and chip as random effects^20^. For the AMDTSS, the Bioconductor package *bacon* was used to adjust for the observed inflation of test statistics. Results from the two cohorts were pooled using fixed-effect metaanalysis^21^. A P-value of 10^−7^ was used to account for multiple testing.

### Associations for methylation sites at genetic loci associated with mammographic density

We annotated the analysed methylation sites to genetic loci associated with mammographic density identified by the largest GWAS to date^4^, using the column ‘UCSC_RefGene_Name’ of the annotation file provided by Illumina. A total of 299 methylation sites were annotated: 4 sites to the *AREG* locus, 76 sites to the *ESR1* locus, 15 sites to the *IGF1* locus, 48 sites to the *LSP1/TNNT3* locus, 60 sites to the *PRDM6* locus, 40 sites to the *SGSM3/MKL1* locus, 17 sites to the *TMEM184B* locus and 39 sites to the *ZNF365* locus. Associations between these 299 sites and mammographic density measures were examined using results from the metaanalysis. The observed number of sites with *P*<0.05 was compared with the expected number using the binomial test.

## Results

None of the studied mammographic density measures was associated with DNA methylation at any site (all *P*>3.4 × 10^−7^ from meta-analysis; Figure 1). No evidence of inflation was found from the meta-analysis: inflation factors were 0.99, 0.97 and 0.96 from the analyses of dense area, non-dense area and percentage dense area, respectively. No genome-wide significant association was found from either of the cohort-specific analyses (all P>1.5 × 10^−7^; Figure 2).

**Figure1.**
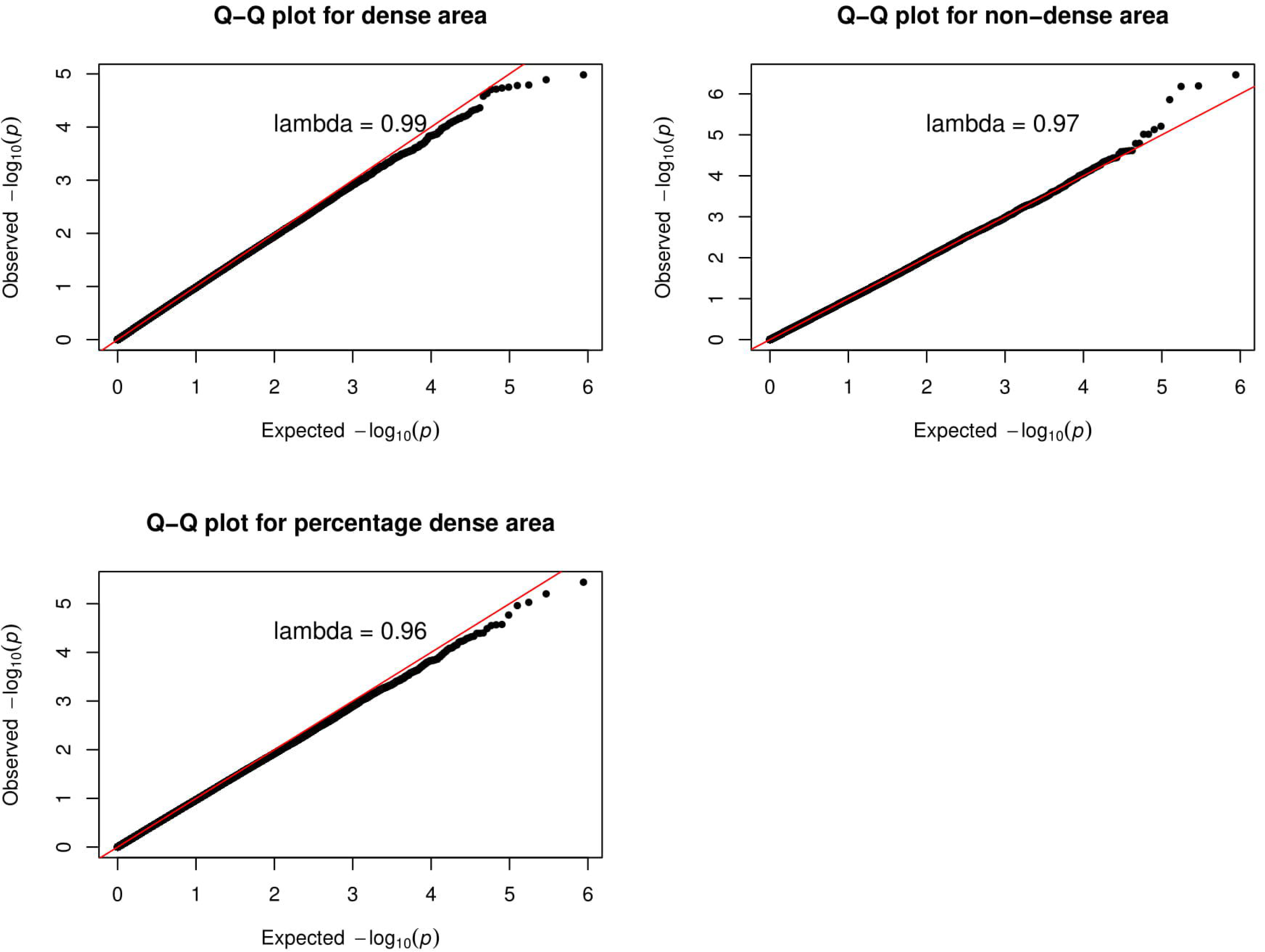
Q-Q plot of the genome-wide association analysis results from the meta-analysis of the two cohorts.

**Figure 2.**
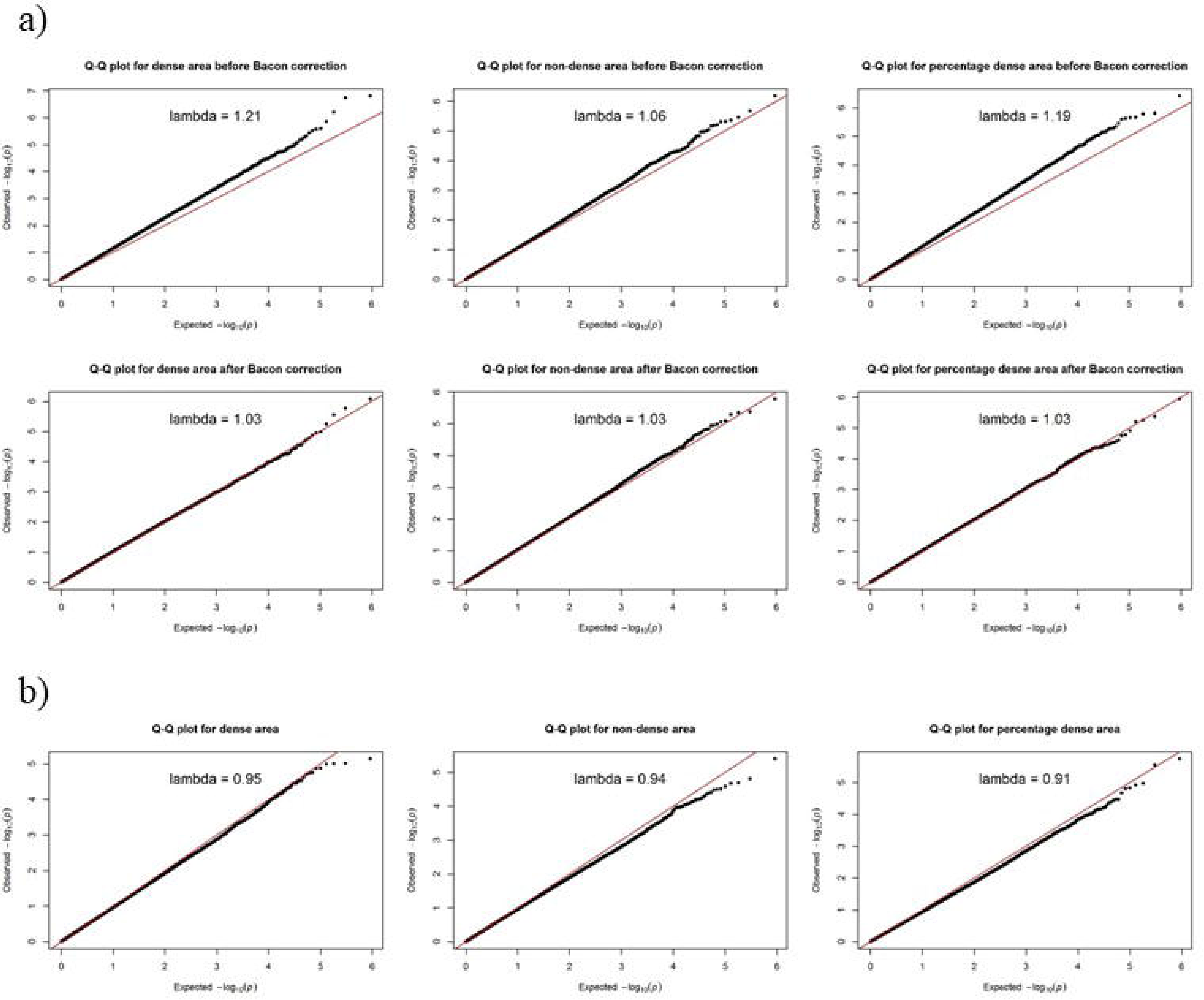
Q-Q plots of the genome-wide association analysis results from the AMDTSS and MCCS a) AMDTSS; b) MCCS.

Of the 299 methylation sites located at the eight genetic loci associated with mammographic density, none was associated with any of the studied mammographic density measures after Bonferroni adjustment for multiple testing (all *P*>0.05/299 = 1.7 × 10^−4^). From the analyses of dense area, non-dense area and percentage dense area, a total of 22, 21 and 21 sites, respectively, had a nominally significant association at *P*<0.05, but this was not significantly greater than expected by chance (all *P*>0.08).

## Discussion

Our genome-wide analysis of data for middle-aged Australian women did not detect associations between blood DNA methylation and conventional mammographic density measures that predict breast cancer risk. No association was found either at any of the loci for which genetic variants were reported to be associated with mammographic density.

Several factors could explain our null results. First, our study included over 1,000 participants, but this might not have been sufficient to detect existed but moderate or weak associations. Second, DNA methylation was measured with the widely used HM450 assay, which covers the promoter regions of 99% RefSeq genes but only a fraction of the genome and might therefore not include sites associated with mammographic density. Third, there could be substantial measurement error in methylation measures obtained from the HM450 assay, which would have weakened the statistical power of our analysis^13, 22^ Fourth, we used the conventional site-by-site analytic approach and a stringent threshold for statistical significance, which might not detect gene- or region-level methylation associated with mammographic density. Fifth, we measured DNA methylation in peripheral blood, not breast tissue, because it is easily accessible and potentially useful for public and clinical health translation^23^. While methylation marks are usually quite stable across tissues, to our knowledge this has not been carefully studied for breast tissue compared with peripheral blood, and not regarding biological relevance to mammographic density. Finally, peripheral blood DNA methylation might not be related to mammographic density at all, or at most trivially, which is indirectly supported by the fact that inconsistent or null results were observed in studies of the relationship between blood DNA methylation and breast cancer risk^8^.

In this study, mammographic density was defined by the conventional brightness threshold using the CUMULUS software and has been referred to as *Cumulus.* New mammographic density measures defined by higher pixel brightness thresholds, *Cirrocumulus* and *Altocumulus*, have been found to predict breast cancer risk better than the *Cumulus* measures, and multivariate analyses suggest that the *Cumulus* measures are only surrogates for the causal aspects of a mammographic image that predict risk^24^. A new breast cancer risk-predicting measure called *Cirrus* has also been developed using machine learning^25^. The *Cirrocumulus*, *Altocumulus* and *Cirrus* measures might be more aetiologically relevant to breast cancer risk. Further research is therefore warranted to investigate associations between DNA methylation and these new mammographic density measures.

Our study has two main strengths. First, to the best of our knowledge, it is the first to investigate the association between DNA methylation and mammographic density. Second, we pooled data from relatively homogenous samples of middle-aged Australian women, for whom DNA methylation and mammographic density were measured using the same protocols.

In summary, no association between blood DNA methylation measured by the HM450 assay and conventional mammographic density measures that predict breast cancer risk was detected by our analysis of middle-aged Australian women.

## Ethics approval and consent to participate

The AMDTSS and MCCS were approved by the Human Research Ethics Committees of the University of Melbourne and Cancer Council Victoria, respectively. All participants provided informed consent.

## Consent for publication

Not applicable.

## Authors’ contributions

Study design: SL, PAD, LB, DRE, MCS, GGG, JLH and RLM. Data collection and preparation: LB, GS, EMW, TLN, JS, DRE, MCS, GGG, JLH and RLM. Data analysis: SL and PAD. Data interpretation: SL, PAD, GS, GGG, MCS, JLH and RLM. Manuscript drafting - SL, PAD, JLH and RLM. All authors contributed to the preparation of and approved the final version of the manuscript.

## Acknowledgements

We would like to thank all women participating in the two cohorts. The AMDTSS was facilitated through access to Twins Research Australia, a national resource supported by a Centre of Research Excellence Grant (ID: 1079102), from the NHMRC. The AMDTSS data analysis was facilitated by Spartan, the High Performance Computer and Cloud hybrid system of the University of Melbourne. SL is a Cancer Council Victoria Postdoctoral Research Fellow. MCS is a NHMRC Senior Research Fellow (APP1061177). JLH is a NHMRC Senior Principal Research Fellow.

